# Improved pure tone sensitivity after simulated hearing loss

**DOI:** 10.1101/2020.05.29.124321

**Authors:** Patrick Krauss

**Affiliations:** Neuroscience Lab, Experimental Otolaryngology, University Hospital Erlangen, Germany; Cognitive Computational Neuroscience Group, Chair of Linguistics, University Erlangen-Nürnberg (FAU), Germany; Department of Otolaryngology, University Medical Center Groningen, The Netherlands

**Keywords:** Hearing loss, tinnitus, Zwicker tone, notched acoustic stimulation, phantom sounds, auditory system, stochastic resonance, ear plugging

## Abstract

Recently, it was proposed that a processing principle called *adaptive stochastic resonance* plays a major role in the auditory system, and serves to maintain optimal sensitivity even to highly variable sound pressure levels. As a side effect, in case of reduced auditory input, such as permanent hearing loss, this mechanism may eventually lead to the perception of phantom sounds like tinnitus or the Zwicker tone illusion. Using computational modeling, the biological plausibility of this processing principle was already demonstrated. Here, we provide empirical results that further support the stochastic resonance model of auditory perception. In particular, Mongolian gerbils were exposed to long-term notched noise, which mimics hearing loss for frequencies within the notch. Remarkably, the animals developed increased sensitivity, i.e. improved hearing thresholds, for the frequency centered within the notch, but nut for frequencies outside the notch. In addition, most animals treated with the new paradigm showed identical behavioral signs of phantom sound perception as animals with acoustic trauma induced tinnitus. In contrast, animals treated with broadband noise as a control condition did not show any significant threshold change, nor behavioral signs of phantom sound perception.

## Introduction

Whether listening to nearly inaudible quiet whispering or enjoying a loud rock concert, the human auditory system has a remarkable ability to adapt to changing sound pressure levels covering several orders of magnitude from the absolute threshold of hearing to the threshold of pain and beyond. Several studies even demonstrate - under certain conditions - the auditory system’s ability to further improve sensitivity even below the absolute threshold of hearing (Wiegrebe et al., 1996; Zeng et al., 2000; Long et al., 2004; Ries, 2007). However, until recently the underlying neural processes remained rather elusive.

In recent studies, we argued that a processing principle called *adaptive stochastic resonance* (Krauss et al., 2017) is exploited by the auditory system in order to continuously maintain optimal sensitivity even to highly variable sound pressure levels and changing statistics of the acoustic environment (Krauss et al., 2016; 2018). The term stochastic resonance refers to a phenomenon, where a signal of arbitrary kind, which is too weak for a certain sensor for being detected, can be made detectable by adding a random signal, i.e. noise, of appropriate intensity to the sensor input (Benzi et al., 1981; Gammaitoni et al, 1998). In the last decades, stochastic resonance has been found in a vast number of different organisms and biological systems (McDonnell et al., 2008; McDonnell & Abbott, 2009). In particular, in neuroscience stochastic resonance helps to explain how nervous systems robustly operate in noise environments (Hänggi, 2002).

According to our model, stochastic resonance is a major processing principle of the auditory system, and takes place in the dorsal cochlear nucleus (Krauss et al., 2016). There, auditory input from the cochlea converges with projections from the somatosensory system (Shore & Zhou, 2006). Since this somatosensory input is largely uncorrelated with the auditory signals, we argued that these somatosensory projections serve as a random signal, i.e. noise, which is necessary for stochastic resonance to work (Krauss et al., 2018). The intensity of the noise is continuously adjusted, depending on the statistics of the auditory input. In case of reduced auditory input for instance, the internal noise would be upregulated, i.e. somatosensory projections dis-inhibited, which results in increased sensitivity by means of stochastic resonance, thereby enhancing information transmission from the cochlea to the auditory system. This assumption is supported by empirical findings that somatosensory projections to the cochlear nucleus are actually upregulated after unilateral deafness (Shore et al., 2008; Dehmel et al., 2012; Zeng et al., 2012).

By constructing a computational model, we further demonstrated the biological plausibility of the proposed processing principle, and demonstrated that stochastic resonance may even improve, i.e. partly restore, speech recognition after hearing loss (Schilling et al., 2020a). The increased internal noise, i.e. upregulated somatosensory input to the auditory system corresponds to the observed permanent increase of spontaneous firing rates within the auditory system in case of chronic hearing loss (Krauss et al., 2016). This neural hyperactivity in turn, was found to be correlated with subjective tinnitus in many studies. Our model provides a unified mechanistic explanation how hearing loss, phantom perceptions like tinnitus and Zwicker tone (Zwicker 1964), and neural hyperactivity are related to each other (Schilling et al., 2020b). Finally, our model predicts that patients with hearing loss and with tinnitus on average have better hearing thresholds than those without tinnitus: tinnitus is the perception of the increased neural noise from the somatosensory system, and this noise helps to increase auditory sensitivity by means of stochastic resonance. By evaluating data from almost 40,000 patients, we could already confirm this prediction (Gollnast et al., 2017).

In order to provide further evidence for the hypothesis that adaptive stochastic resonance plays a major role in the auditory system and especially in phantom sound perception, we developed a novel animal experimental paradigm: *simulated hearing loss through long-term noise exposure with notched noise*.

In nature, the standard acoustic environment, i.e. the temporally averaged frequency power spectrum, is broad and flat, i.e. all frequencies have statistically similar mean sound pressure levels - at least on time scales of days to weeks, and at least in the frequency range best perceivable by animals and humans (Schaette and Kempter, 2006). Therefore, it seems possible to simulate a certain hearing loss by exposure with sound or noise where the mean (i.e. temporally averaged) power of the frequency band of desired hearing loss is decreased (= dipped) compared to adjacent frequencies. Hence, the otherwise flat power spectrum has a dip that mimics hearing loss. From the perspective of the auditory system, it should make no difference whether certain frequency channels receive less input due to cochlear damage or due to changed statistics of the acoustic environment. After the end of long-term noise exposure, animals’ acoustic environment has a flat audio power spectrum on longer time scales again which, analogously to the above argument, corresponds to a complete restoration of hearing. Thus, the post-noise exposure phase of the paradigm corresponds to a complete restoration, or simulated treatment with an ideal hearing aid, respectively.

## Results

Eleven animals (Mongolian gerbils) have been treated with the new paradigm (spectral dipped noise), and additional six animals with the control paradigm (spectral flat noise). The relative threshold changes during long treatment are summarized in Figure 1. Animals treated with spectral dipped noise showed a significant (p=0.022) mean decrease (= improvement) of hearing thresholds of 5 dB for 2 kHz, exactly where the dip was centered. In contrast, hearing thresholds for 1 and 4 kHz (where the spectrum was flat) did not change significantly (Figure 1, left panel). The control group stimulated with spectral flat noise showed no significant shift in any tested frequency (Figure 1, right panel). These results are perfectly in line with the hypothesis that the auditory system exploits stochastic resonance to compensate for changing input statistics.

**Figure 1:**
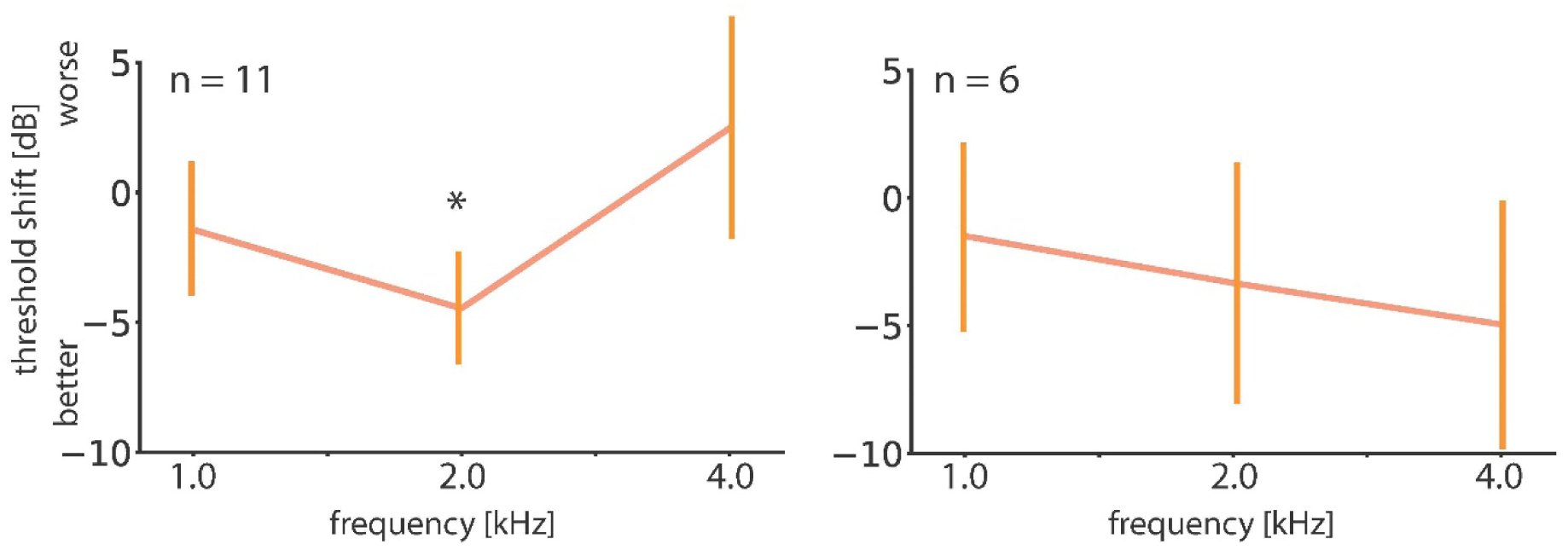
Animals treated with spectral dipped noise (n=11) showed a significant (p=0.022) mean decrease (= improvement) of hearing thresholds of 5 dB for 2 kHz, exactly where the dip was centered. The hearing thresholds for 1 and 4 kHz (where the spectrum was flat) did not change significantly (left panel). In contrast, the control group stimulated with spectral flat noise (n=6) showed no significant shift in any tested frequency (right panel).

In addition, most animals (ten of eleven) treated with the new paradigm showed identical behavioral signs of phantom sound perception as animals with acoustic trauma induced tinnitus, whereby the strongest acute effects occurred at the frequency where the spectral dip was centered (2 kHz). In the control group, no behavioral signs of phantom sound perception was observed. Again, this supports our stochastic resonance model of tinnitus development. Only one animal did not show any behavioral signs of tinnitus, similar to the observed ratio of tinnitus development after noise trauma.

After the end of long-term notched noise exposure which corresponds to subsequent simulated treatment with a perfect hearing aid, one group of animals (n=4) showed a complete removal of behavioral signs of tinnitus, whereas another group did not respond to the end of the treatment. In those animals (n=6) the tinnitus perception became chronical and could be observed even two weeks after the end of notched noise exposure. Remarkably, these results correspond to what is known from tinnitus patients treated with hearing aids or cochlea implants. In one group of patients, the tinnitus perception is reduced or even completely removed, whereas another group does not respond to the treatment (Henry et al., 2015; Jalilvand et al., 2015; Mazurek et al., 2017).

## Discussion

We presented a novel experimental paradigm to simulate transient hearing loss, which in turn induced improvement of hearing thresholds and perception of phantom sounds, both in line with our hypothesis that stochastic resonance plays a major role in the auditory system. The converging evidence from numerous empirical (for an overview see Krauss et al., 2018 and Schilling et al., 2020b) and theoretical studies (Krauss et al., 2016; 2017; Schilling et al., 2020a) indicates that the auditory system actually exploits stochastic resonance and actively tunes the intensity of required noise by adjusting the somatosensory input to the dorsal cochlear nucleus. By that, stochastic resonance contributes crucially to the auditory system’s ability to adapt to changing sound pressure levels.

In terms of time scale, the induced phantom sounds are at an intermediate level between chronic lifelong tinnitus on the one end and the only seconds lasting Zwicker tone percepts (Zwicker 1964) on the other end of the spectrum. Within the stochastic resonance framework of auditory processing, all these phantom sounds are caused by the same mechanism and do just occur on different time scales (Schilling et al., 2020b).

Furthermore, the presented results are similar to ear plugging studies, where healthy human subjects are provided with ear plugs for two weeks, mimicking hearing loss. These subjects perceive a transient tinnitus after removal of ear plugs (Schaette et al., 2012; Fournier et al., 2014). In contrast to ear plugging, where mainly higher frequencies are attenuated, the long-term noise exposure paradigm has the advantage that both, spectral location and shape of simulated hearing loss are fully controllable.

In follow-up studies, it might be interesting to systematically compare different shapes of hearing loss, e.g. high-frequency hearing loss versus low-frequency hearing loss. Furthermore, the impact of other kinds of noise (e.g. babble noise or correlated noise) on hearing threshold changes and induction of phantom sounds, may be investigated.

Finally, we note that our paradigm is an important step towards reducing the health burden on laboratory animals, since instead of applying a permanent noise trauma in order to induce tinnitus, the hearing loss can now be simulated.

## Methods

### Animals and ethics statement

Rodents (Mongolian gerbils (*Meriones unguiculatus*) were housed in standard animal racks (Bio A.S. Vent Light, Zoonlab GmbH, Castrop-Rauxel, Germany) in groups of 2 animals per cage with free access to water and food at 20 to 24°C room temperature under 12/12 h dark/light cycle. The use and care of animals was approved by the state of Bavaria (Regierungspräsidium Mittelfranken, Ansbach, Germany; AZ: 54-2532.1-02/13 and 54-2532.1-42/13). Gerbils were purchased from Charles River Laboratories Inc. (Sulzfeld, Germany).

### Generation of noise stimuli

Starting from white noise with flat frequency spectrum, dipped noise has been generated with equalizer software. The dip has a spectral width of an octave centered on a frequency of 2 kHz. At the center frequency the power-difference to the non-dipped (=flat) spectrum is maximal (20 dB) and decreases sigmoidally (in order to avoid sharp edges in the spectrum) with increasing distance to the center. At the borders (1.4 and 2.8 kHz) the power-difference is zero, i.e. the power is identical to the rest of the power spectrum. For the control group non-dipped (= flat) white noise was used.

### Long-term noise exposure and experimental protocol

Exposure to dipped/flat noise at low sound pressure levels (50dB SPL) took two weeks for each animal. The notched noise had a dipped spectral envelope. This new paradigm simulates both hearing loss (dip in frequency spectrum) and potential subsequent treatment with an ideal hearing aid or therapy that fully restores hearing abilities. Hearing thresholds for frequencies of 1, 2 and 4 kHz were measured using auditory brainstem responses in all animals before and immediately after long-term noise exposure. The GPIAS paradigm (Turner et al., 2006; Schilling et al., 2017) was used to assess the potential existence of a tinnitus percept. Animals were tested before (pre), immediately after (acute), one week and two weeks after long-term noise exposure.

### Assessment of Hearing Thresholds

In order to assess the animal’s hearing thresholds, auditory brainstem responses (ABR) were recorded using a custom made setup (Schilling et al., 2019). Pure tone stimuli of different frequencies (1.0, 2.0 and 4.0 kHz) were generated by a custom-made python program and presented at different, pseudo-randomized intensities ranging from 90 to 0 dB SPL in 5 dB steps. Stimulation was free-field to the measured ear (ipsi-lateral) at a time via a speaker (Sinus Live NEO) corrected for its frequency transfer function to be flat within +/-1 dB at a distance of approximately 30 mm from the animal’s pinna while the contra-lateral ear was tamped with an ear plug. To compensate for speaker artifacts stimuli were presented as double trials consisting of two 6 ms stimuli (including 2 ms sine square rise and fall ramps) of the same amplitude but opposite phase, separated by 100 ms of silence. 120 double trials of each combination of intensity and frequency were presented pseudo-randomly at an inter-stimulus interval of 500 ms.

For the measurements the Mongolian gerbils were kept under deep anesthesia. Anesthesia was induced by an initial dose of 0.3 ml of a ketamine-xylacin-mixture (mixture of ketamine hydrochloride: 96 mg/kg BW; xylacin hydrochloride: 4mg/kg BW; atropine sulfate: 1 mg/kg BW), and maintained by continuous application of that mixture at a rate of 0.2 to 0.3 ml/h by a syringe pump. As has been demonstrated previously, such ketamin-xylazine anesthesia has only little effect on ABR signals compared to awake animals (Smith and Mills 1989)

During measurements, animals were placed on a heating pad at 37°C. Data were recorded using three silver wire electrodes positioned subcutaneously, one for grounding at the back of the animals, one reference electrode at the forehead and the measuring electrode infra-auricular overlying the bulla. The potential difference between the reference and measuring electrode was amplified by a low noise amplifier (JHM NeuroAmp 401, J. Helbig Messtechnik, Mainaschaff, Germany; amplification 10.000; bandpass filter 400 Hz to 2000 Hz and 50 Hz notch filter). Note that for further analysis the amplified signal was used, that is, amplitudes are given in mV whereas the actual neuronal signals were in µV range. The output signal of the amplifier was digitalized and recorded by an analog-digital converter card (National Instruments Corporation, Austin, TX, USA) with a sampling rate of 20 kHz and synchronized with the stimulation via the trigger signal of the stimulation computer. Raw data of 120 double trials per sound level for one stimulus frequency were averaged. Finally, these averaged responses of the two single, phase inverted stimuli within one double trial were averaged to eliminate stimulus artifacts. From these averaged, artifact-corrected data the root mean square (RMS) values within a time window from 0 to 10 ms after stimulus onset were calculated to obtain a measure of auditory brainstem response amplitude for each stimulus intensity presented.

Hearing thresholds were estimated using the method described in Schilling et al., 2019.

### Phantom sound assessment (GPIAS)

Putative perception of phantom sounds was assessed with the gap pre-pulse inhibition of the acoustic startle reflex (GPIAS) paradigm (Turner et al., 2006) using a custom made setup (Gerum et al., 2019). Date were analyzed using the method described in Schilling et al., 2017.

## Acknowledgements

This work was funded by the Deutsche Forschungsgemeinschaft (DFG, German Research Foundation): grant KR5148/2-1 to PK, project number 436456810

I thank Franziska Schilmeier and Olaf Wendler for technical support, and Holger Schulze for providing me access to the required setups.

## References

Benzi, R., Sutera, A., & Vulpiani, A. (1981). The mechanism of stochastic resonance. Journal of Physics A: mathematical and general, 14(11), L453.

Dehmel, S., Pradhan, S., Koehler, S., Bledsoe, S., & Shore, S. (2012). Noise overexposure alters long-term somatosensory-auditory processing in the dorsal cochlear nucleus—possible basis for tinnitus-related hyperactivity?. Journal of Neuroscience, 32(5), 1660–1671.

Fournier, P., Schonwiesner, M., and Hebert, S. (2014) Loudness modulation after transient and permanent hearing loss: implications for tinnitus and hyperacusis. Neuroscience 283:64–77.

Gammaitoni, L., Hänggi, P., Jung, P., & Marchesoni, F. (1998). Stochastic resonance. Reviews of modern physics, 70(1), 223.

Gerum, R. C., Rahlfs, H., Streb, M., Krauss, P., Grimm, J., Metzner, C., … & Schilling, A. (2019). Open (G) PIAS: An open-source solution for the construction of a high-precision acoustic startle response setup for tinnitus screening and threshold estimation in rodents. Frontiers in Behavioral Neuroscience, 13, 140.

Gollnast, D., Tziridis, K., Krauss, P., Schilling, A., Hoppe, U., & Schulze, H. (2017). Analysis of audiometric differences of patients with and without tinnitus in a large clinical database. Frontiers in Neurology. Doi: 10.3389/fneur.2017.00031

Hänggi, P. (2002). Stochastic resonance in biology how noise can enhance detection of weak signals and help improve biological information processing. ChemPhysChem, 3(3), 285–290.

Henry, J. A., Frederick, M., Sell, S., Griest, S., & Abrams, H. (2015). Validation of a novel combination hearing aid and tinnitus therapy device. Ear and Hearing, 36(1), 42–52.

Jalilvand, H., Pourbakht, A., & Haghani, H. (2015). Hearing aid or tinnitus masker: which one is the best treatment for blast-induced tinnitus? The results of a long-term study on 974 patients. Audiology and Neurotology, 20(3), 195–201.

Krauss, P., Tziridis, K., Metzner, C., Schilling, A., Hoppe, U., & Schulze, H. (2016). Stochastic resonance controlled upregulation of internal noise after hearing loss as a putative cause of tinnitus-related neuronal hyperactivity. Frontiers in Neuroscience. Doi: 10.3389/fnins.2016.00597

Krauss, P., Metzner, C., Schilling, A., Schütz, C., Tziridis, K., Fabry, B., & Schulze, H. (2017). Adaptive stochastic resonance for unknown and variable input signals. Scientific Reports. Doi: 10.1038/s41598-017-02644-w

Krauss, P., Tziridis, K., Schilling, A., & Schulze, H. (2018). Cross-modal stochastic resonance as a universal principle to enhance sensory processing. Frontiers in neuroscience, 12, 578.

Long, Z. C., Shao, F., Zhang, Y. P., and Qin, Y. G. (2004). Noise-enhanced hearing sensitivity. Physics Letters A, 323(5), 434–438.

Mazurek, B., Szczepek, A. J., & Brueggemann, P. (2017). Tinnitus–Klinik und Therapie. Laryngo-rhino-otologie, 96(01), 47–59.

McDonnell, M. D., Stocks, N. G., Pearce, C. E., & Abbott, D. (2008). Stochastic resonance. Stochastic Resonance, by Mark D. McDonnell, Nigel G. Stocks, Charles EM Pearce, Derek Abbott, Cambridge, UK: Cambridge University Press, 2008.

McDonnell, M. D., & Abbott, D. (2009). What is stochastic resonance? Definitions, misconceptions, debates, and its relevance to biology. PLoS computational biology, 5(5).

Ries, D.T. (2007). The influence of noise type and level upon stochastic resonance in human audition. Hearing Research 228, 136–143

Schaette, R., and Kempter, R. (2006) Development of tinnitus-related neuronal hyperactivity through homeostatic plasticity after hearing loss: a computational model. European Journal of Neuroscience, Vol. 23, 3124–3138

Schaette, R., Turtle, C., and Munro, K.J. (2012) Reversible induction of phantom auditory sensations through simulated unilateral hearing loss. PloS one 7:e35238.

Schilling, A., Krauss, P., Gerum, R., Metzner, C., Tziridis, K., & Schulze, H. (2017). A new statistical approach for the evaluation of gap-prepulse inhibition of the acoustic startle reflex (GPIAS) for tinnitus assessment. Frontiers in Behavioral Neuroscience. Doi: 10.3389/fnbeh.2017.00198

Schilling, A., Gerum, R. C., Krauss, P., Metzner, C., Tziridis, K., & Schulze, H. (2019). Objective estimation of sensory thresholds based on neurophysiological parameters. Frontiers in neuroscience, 13, 481.

Schilling, A., Gerum, R., Zankl, A., Schulze, H., Metzner, C., & Krauss, P. (2020a). Intrinsic noise improves speech recognition in a computational model of the auditory pathway. bioRxiv.

Schilling, A., Tziridis, K., Schulze, H., & Krauss, P. (2020b). The Stochastic Resonance model of auditory perception: A unified explanation of tinnitus development, Zwicker tone illusion, and residual inhibition. bioRxiv.

Shore, S. E., & Zhou, J. (2006). Somatosensory influence on the cochlear nucleus and beyond. Hearing research, 216, 90–99.

Shore, S. E., Koehler, S., Oldakowski, M., Hughes, L. F., & Syed, S. (2008). Dorsal cochlear nucleus responses to somatosensory stimulation are enhanced after noise-induced hearing loss. European Journal of Neuroscience, 27(1), 155–168.

Smith, D. I., & Mills, J. H. (1989). Anesthesia effects: auditory brain-stem response. Electroencephalography and clinical neurophysiology, 72(5), 422–428.

Turner JG, Brozoski TJ, Bauer CA, Parrish JL, Myers K, Hughes LF, Caspary DM (2006) Gap detection deficits in rats with tinnitus: a potential novel screening tool. Behav Neurosci 120:188–195.

Wiegrebe, L., Koessl, M., and Schmidt, S. (1996) Auditory enhancement at the absolute threshold of hearing and its relationship to the Zwicker tone. Hearing Research 100, 171–180.

Zeng, F.G., Fu, Q.J., and Morse, R. (2000). Human hearing enhanced by noise. Brain Research 869, 251–255.

Zeng, C., Yang, Z., Shreve, L., Bledsoe, S., & Shore, S. (2012). Somatosensory projections to cochlear nucleus are upregulated after unilateral deafness. Journal of Neuroscience, 32(45), 15791–15801.

Zwicker, E. (1964). “Negative afterimage” in hearing. The Journal of the Acoustical Society of America, 36(12), 2413–2415.

